# More than half of annotated human miRNAs are never expressed at levels sufficient for biological function

**DOI:** 10.1101/2024.12.05.626958

**Authors:** Saba Ataei Kachooei, Julie M Bracken, Katherine A Pillman, Philip A Gregory, Cameron P Bracken

## Abstract

MicroRNAs (miRNAs) are widely studied for their role in post-transcriptional gene regulation, often using exogenous overexpression systems to reveal their functions. However, such approaches may not accurately reflect endogenous miRNA activity due to the substantially higher expression levels achieved experimentally. To address this, we sought to determine the minimal endogenous expression threshold required for a miRNA to exert biologically significant effects. By comparing these experimentally determined expression thresholds with small RNA sequencing datasets comprising hundreds of cell lines and tens of thousands of tissue samples, we found that more than half of all annotated miRNAs are never expressed at levels sufficient to be biologically relevant. This calls into question the conclusions of thousands of studies reporting functions for these lowly expressed miRNAs, whose results are likely attributable to artificial overexpression rather than physiological activity. Our study highlights the need for more rigorous evaluation of miRNA functionality in their native context, and provides further support to arguments that the size of the functional human “microRNAome” is far smaller than some estimates of miRNA numbers based upon small RNA sequencing data.

## Introduction

MicroRNAs (miRNAs) act as the target recognition component of a larger ribonucleoprotein complex called RISC (RNA-induced silencing complex), where in a sequence specific manner, they act to bring target mRNAs to RISC via complementary base pairing. RISC then suppresses the target gene through a combination of mRNA destabilisation and translational repression (1, 2). Initially discovered in *Caenorhabditis elegans* in 1993 (3, 4), interest in the field rapidly grew with the subsequent discovery of miRNAs in both plants and mammals. Mechanisms of miRNA biogenesis and function were then quickly uncovered, with each newly discovered miRNA annotated and placed into databases from which miRbase emerged as the primary resource (5).

Perhaps surprisingly however, given the miRNA field spans ∼200k publications, the number of genuine miRNAs encoded within the human genome has remained contentious. Some estimates derived from the analysis of extensive sequencing data has placed the number of human miRNAs at over 5000 (6), whilst others place the number at <600 (7, 8). The contention lies with separating genuine, functional miRNAs from the RNA degradome, and to what extent non-canonical features should be used as exclusionary factors. For example, to what degree should one require high 5’-homogeneity, high “in-cluster ratio” (the proportion of local reads mapping to a specific miRNA) and the presence of sequences mapping to the other arm of a hairpin structure from which miRNAs are canonically produced (8)? These questions are especially difficult given there are non-canonical pathways that can give rise to functional miRNAs (9-15).

Beyond the question of what features of small RNAs should be regarded as exclusionary, there is also a fundamental question of how much miRNA needs be present to be biologically meaningful? MiRNA activity is dose-dependent, but since the use of miRNA mimics leads to supra-physiological levels of expression, and the displacement of endogenous miRNAs from AGO, their use may not be representative of authentic miRNA function (16). Further, if a contentiously annotated miRNA is expressed at levels insufficient to be biologically meaningful, the status of the RNA itself is moot.

To address this, we sought to investigate the minimal level of endogenous expression that is required to warrant consideration of miRNA function. To do so, reporters were constructed for multiple miRNAs and their activity examined in three cell lines that naturally possess differing levels of the selected miRNAs. By cross-referencing reporter activity against endogenous miRNA expression as determined by RNA-seq, one can estimate the minimal amount of miRNA that needs to be present for detectable activity. In general agreement with previous work (17, 18), even when we use reporters that are fully complementary to the miRNA and thus, should be far more efficiently repressed than any endogenous target in cells, we note miRNA activity is never detectable at less than 200 counts per million (cpm, as determined by small RNA-seq). At levels greater than 1000 cpm the activity of most miRNAs is discernible, though higher expression does not necessarily always correlate with stronger repressive activity.

We then use this information to interrogate small RNA sequencing data and find that from the 1743 different miRNA families annotated in miRBase, 1042 are never expressed at a level (>100 cpm) at which activity would be expected in any of the >300 cell lines and >10,000 tissues examined. These public data are drawn from hundreds of studies, almost 200 different tissue sites and a variety of normal and pathological conditions (19), meaning it is unlikely we are simply missing a context in which specific small RNAs are highly expressed and functional. Despite this seeming incapacity to exert biologically meaningful functions, regardless of whether they are genuine miRNAs (incorporated into RISC) or merely products of the degradome, the lowly expressed miRNAs are the subject of >4,500 papers featuring the name of the miRNA in the title, and >10,000 papers where the miRNA features in the abstract. This work highlights the importance of considering endogenous miRNA expression, and calls into question the significance of thousands of studies whose conclusions we expect are solely attributable to supraphysiological expression.

## Materials and Methods

### Cell culture

HEK-293T, HeLa and MDA-MB-231 cells were cultured in DMEM (Dulbecco’s Modified Eagle Medium-Gibco) supplemented with 10% fetal calf serum (FCS; Cytiva, #SH30071.03). All cell lines were incubated at 37°C in a humidified atmosphere containing 5% CO_2_, with mycoplasma testing performed regularly.

### Reporters construct design and cloning

The PsiCHECK2 dual luciferase reporters (Promega, #C8021) were designed to incorporate a single copy of a target site that perfectly complements small RNA sequences from miRBase (5). Oligonucleotides (IDT technologies) were created with overhangs for XhoI (New England BioLabs, #R0146S) and NotI-HF (New England BioLabs, #R3189S), which were annealed and ligated (NewEngland BioLabs, # M0202S) into psiCHECK2 using the same restriction sites.

### Luciferase reporter assays

HeLa and HEK-293T cells were plated in 96-well plates at densities of 10,000 and 20,000 cells per well, respectively, while MDA-MB-231 cells were seeded in 24-well plates at a density of 100,000 cells per well 1 day before transfection. Prior to transfection, the media was changed, and the cells were co-transfected with 5ng of miRNA psiCHECK2 reporter constructs using Lipofectamine™ 2000 Transfection Reagent (Invitrogen, #11668019), following the manufacturer’s instructions. The media was changed 6 hours after transfection. After 48 hours, luciferase activity assay was conducted using the Dual-Luciferase® Reporter Assay System (Promega, #E1980) on a GloMax®-Multi Detection System (Promega), in accordance with the manufacturer’s guidelines. The relative luciferase activity was determined by calculating the ratio of Renilla to Firefly luciferase, with the Firefly luciferase gene expressed from the same vector serving as an internal control, while cells transfected with the empty psiCHECK2 vector acted as a control group. All reporter assays were conducted in four technical replicates for each experiment, which was repeated independently three times.

### Small RNAseq

For each cell line RNA was extracted according to Trizol Reagent (Ambion, #15596018) manufacturer instructions. The quantity and of the RNA samples were checked using nanodrop Qubit HS RNA (Invitrogen, #Q32852) and Bioanalyser small RNA assay (Agilent, #5067-1548). Libraries were generated using 1 ug of the total RNA via QIAseq miRNA kit (Qiagen, # 331502) with 14 cycles of amplification. Amplified barcoded libraries were then size selected using auto gel-purification Pippin prep 3% agarose (SAGE science) which targets ranges from 100-250bp. Libraries (size between 180-190bp) were then confirmed by Qubit HS DNA and Bioanalyzer HS DNA assay for size and concentration. The libraries were then pooled together in equimolar amounts and sequenced using an Illumina MiSeq Reagent Kit v3 kit in accordance with the manufacturer’s guidelines. Methodological details for the AGO:RNA co-immunoprecipitation (Supplementary Figure 1) and the growth of HMLE and mesHMLE cell lines are fully detailed in (Orang et al, Under Review).

### Data sources for bioinformatic assessment

Small RNA sequencing data was obtained from miTED (19). Tissue-specific 5p/3p miRNA expression data (Supplementary Figure 2) was obtained from miRSwitch (20).

## Results

### Constructing sensitive reporters of endogenous miRNA activity

Our goal is to establish the minimal level of expression at which an individual miRNA might be functional, then interrogate large public sequencing repositories to assess how often different miRNAs meet these thresholds. Therefore, we first sought to establish a highly sensitive system to report on endogenous miRNA activity. Although mammalian miRNAs rarely interact with their targets with full sequence complementarity, we rationalised fully complementary reporters should be the most sensitive as the extensive miRNA:target interaction interface leads to stronger binding and the unmasking of AGO2’s endogenous slicer activity (directly cleaving the target mRNA, (21-23). To confirm this strategy, both full complementary and seed-only (endogenous-like) binding sites were engineered downstream of a Renilla luciferase reporter for two highly expressed miRNAs; miR-21-5p (the 3^rd^ highest expressed miRNA in HeLa cells at 62k cpm) and miR-29-3p (the 12^th^ highest expressed miRNA at 25k cpm). In both cases, activity of the perfect reporter was vastly decreased relative to both unmodified Renilla luciferase and the seed-only reporter. In the case of miR-21-5p, exogenous miR-21-5p has no additional effect, presumably as cellular AGO was already sufficiently loaded with miR-21 to maximally suppress activity of the introduced reporter. In the case of miR-29a-3p, additional miRNA was able to exert further repression (Figure 1A). These results indicate fully complementary reporters are far more sensitive than seed-only reporters and that at least for highly expressed miRNAs, luciferase/reporter activity is strongly depleted by endogenous miRNA levels. As the degree to which a target can be repressed is a balance between the level of target and repressor, we sought to establish optimal levels of reporter transfection such that the amount of reporter yielded reproduceable results, but was minimised as much as possible to ensure maximal sensitivity (Figure 1B). On the basis of these data, 5ng psiCHECK2 was selected. Additional fully complementary reporters were then constructed to measure the activity of miRNAs ranging from those with little to no endogenous expression, through to those which are highly expressed.

**Figure 1.**
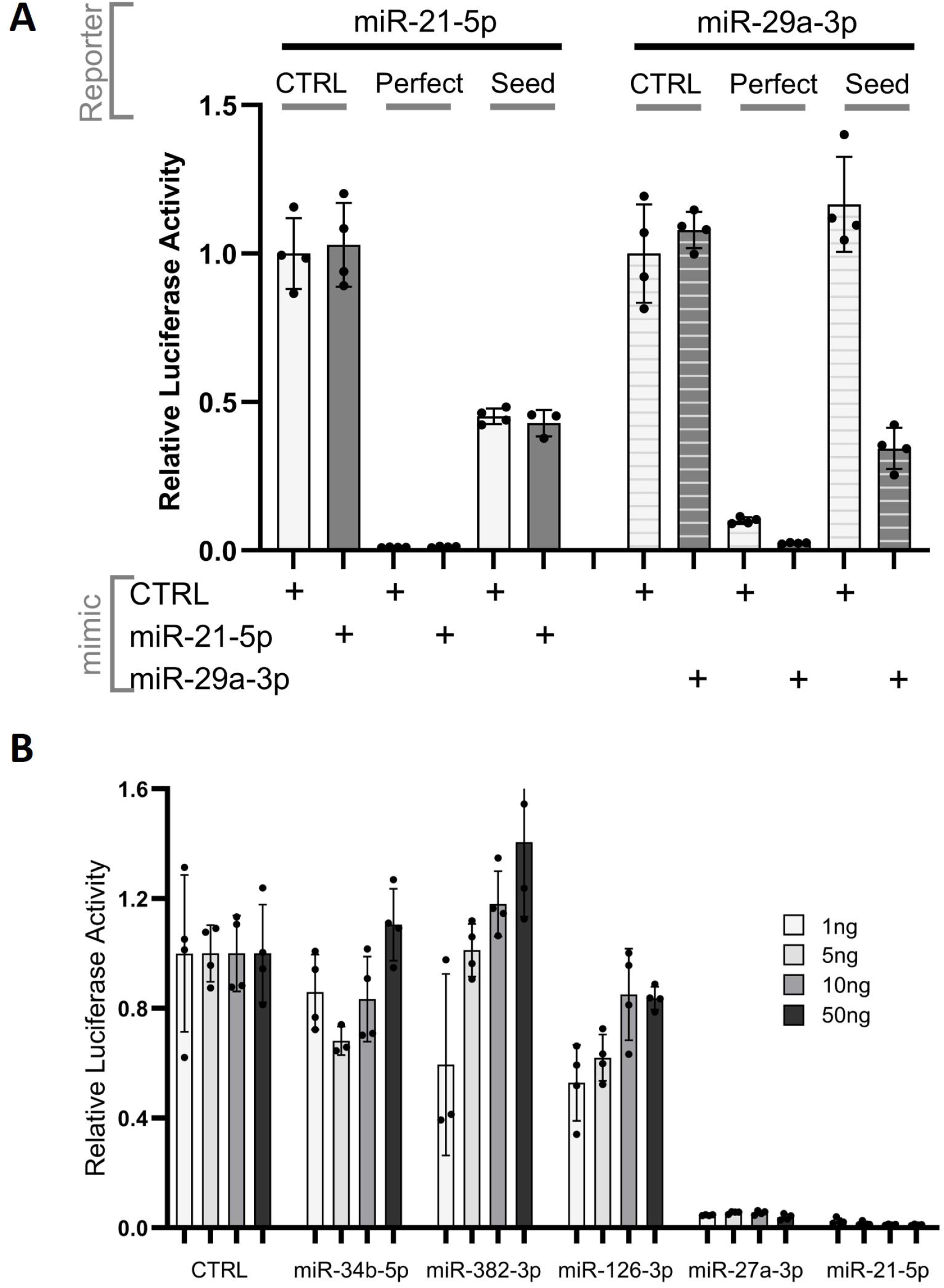
Constructing sensitive reporters of endogenous miRNA activity. A. Renilla luciferase reporters were constructed in which the reporter was either unmodified (CTRL), or where sites for miR-21-5p or miR-29a-3p were introduced into the Renilla luciferase 3’UTR that are either fully complementary across the entire length of the 21nt binding interface (“perfect”) or complementary only across an 8nt seed-site (“seed”, nt 2-9 inclusive relative to the 5’ end of the miRNA). Dual Renilla luciferase reporter assays were then conducted, co-transfecting the Renilla luciferase expression plasmid along with negative control miRNA mimic, or mimics to miR-21 and miR-29a. B. Renilla luciferase vectors constructed to report on the activities of the miRNAs indicated were transfected into HeLa cells at 1, 5, 10, 50 ng.

### Endogenous miRNA activity is undetectable for miRNAs expressed below several hundred counts per million

Using a dual-luciferase reporter assay, the activity of 20 reporter genes engineered to be responsive to the presence of the miRNAs indicated were quantitatively assessed in three cell lines: HeLa (Figure 2A), MDA-MB-231 (Figure 2B), and HEK-293T (Figure 2C). There is a clear trend that miRNA reporter activity decreases as endogenous expression of the corresponding miRNA increases, indicating reporters provide a proportional readout of miRNA presence. However, reporter systems display intrinsic variability, on account of such factors as the presence of cryptic sites for miRNAs other than that to which they were designed, or the creation of binding sites for other RNA-binding proteins of which there are many that associate with 3’UTRs (24). For this reason, we see variation in reporter activity, even in some cases where a reporter has been designed to a miRNA which is not expressed and for which there are no other obvious miRNA binding candidates. For this reason, the “group CTRL” bar has been added which is the summation of measurements for all reporters designed against miRNAs for which detectable endogenous activity is expected to be negligible (<20 cpm). The shaded horizontal yellow bar indicates the upper and lower bounds of activity for reporters for miRNAs expressed at <20 cpm. Statistical significance was then calculated for each reporter against this group control. This is indicated in the corresponding table, as are read counts of the miRNA to which the reporter was designed (“exact”) and read counts of other miRNA family members that share the same seed site (“seed”) and thus, would be expected to target the reporter, but to do so less efficiently (in a non-siRNA like manner). This is why for example, the miR-99a-5p reporter is so strongly suppressed in MDA-MB-231 cells, as although miR-99a-5p expression is very low, its seed-matched counterpart miR-99b-5p, is present at far higher levels.

**Figure 2.**
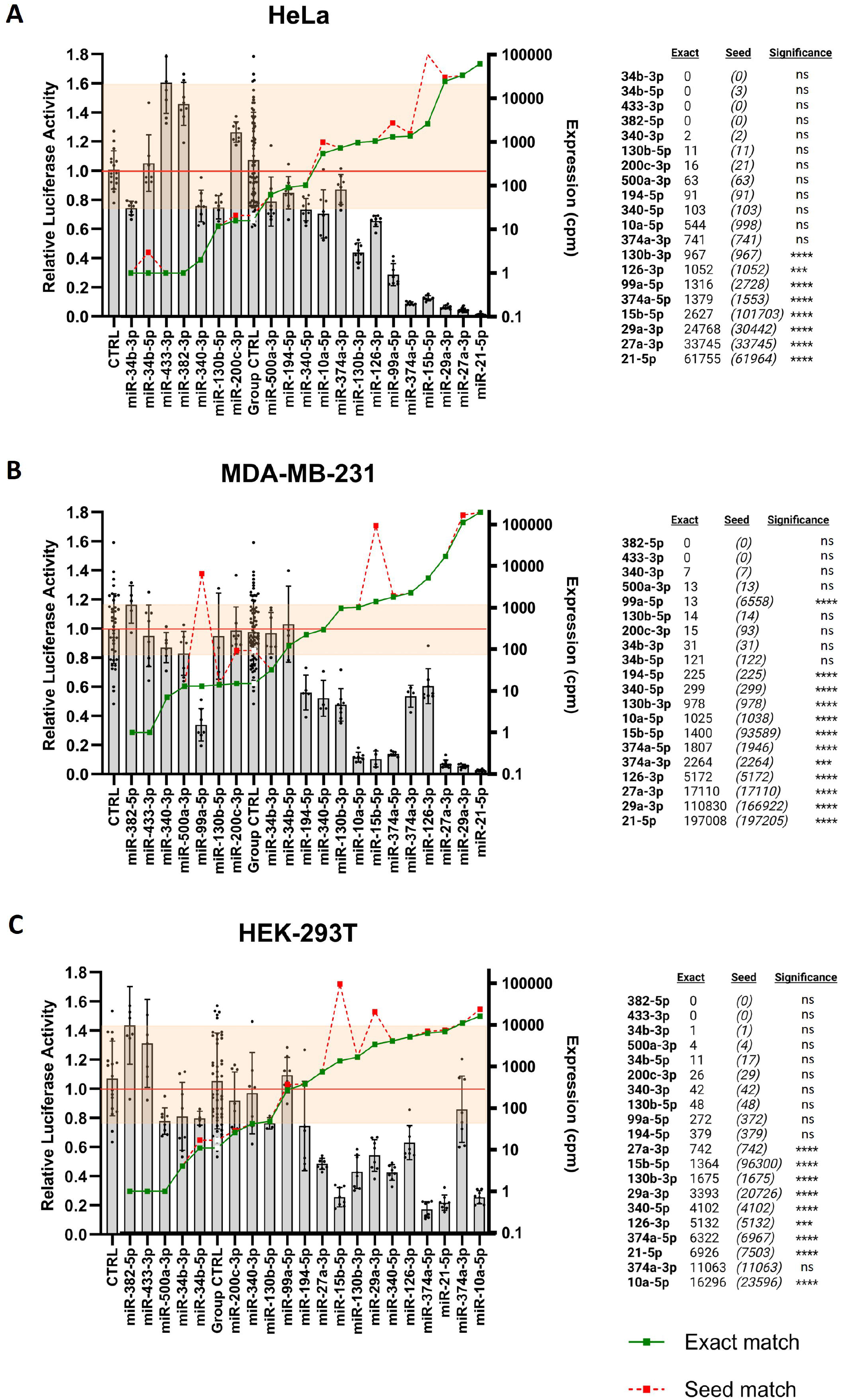
Endogenous miRNA activity is undetectable below 200 cpm. Dual luciferase reporter assays were conducted in A) HeLa, B) MDA-MB-231 and C) HEK-293T cells in which Renilla luciferase reporters that were perfectly complementary to the miRNAs indicated had been transfected. Relative luciferase activity is shown on the left axis, sequencing reads for the miRNA in question on the right (green line). The dotted red line indicates the sum expression of all miRNAs whose seed site would be predicted to also target the reporter. Significance is calculated by t-test, * p=<0.05; ** p=<0.01; *** p=<0.001; **** p=<0.0001. “NS” = not significant

In all cases where a miRNA is expressed above 1000 cpm, statistically significant repression was detected, with the exception of miR-374a-3p in HEK293T cells which, like miR-126-3p, is consistently less active than one would expect solely on the basis of expression. This highlights that not all highly expressed miRNAs produce equivalent functional repression, which may be influenced by factors such the number of competing target sites in other transcripts (25, 26) or AGO-binding affinity. Consistent with this, in data derived from two separate cell lines (mammary epithelial HMLE cells and their mesenchymal “mesHMLE” derivative created by prolonged exposure to TGF-β, (27)), the relative efficiency with which miR-374a-3p is co-precipitated with AGO relative to its expression level detected in whole cell sequencing, was markedly lower than the other miRNAs that exerted suppressive effects on their respective reporters (Supplementary Figure 1).

In no instance was statistically significant repression detected for any reporter designed against a miRNA expressed at less than 200 cpm. The capacity to detect the repressive activities of miRNAs expressed between 200 and 1000 cpm was variable. These findings are in strong agreement with a previous study, where 80% of miRNAs expressed at >1000 cpm showed suppressive activity in a fluorescent-based reporter screen, whilst repression was only detected for <2% of miRNAs expressed at <100 cpm, which the authors speculated may have been false-positives as no repressive activities were detected for any miRNA expressed below this range in a second cell line (18). Full miRNA sequencing data extracted from miTED (19) are provided in Supplementary Table 1.

### Many miRNAs are never expressed at a sufficient level for endogenous functionality

Our data, and others (17, 18), clearly show that miRNAs expressed below 100 cpm have no detectable effect, even on potential targets that have perfect complementarity. Consequently, we examined public small-RNA sequencing data to see how often miRNAs fail to meet this minimum threshold of expression. Figure 3 presents an analysis of 2,656 miRNAs (representing 1743 distinct loci) sourced from miRBase, with varying levels of annotation—some miRNAs have designated 3p or 5p arms, while others lack specific arm assignments. The maximal level of expression for each miRNA across 333 cell lines (Figure 3A) and 10,740 tissue samples (Figure 3B, representing 176 different tissues from both normal and pathologic states) is shown. Importantly, 1,425 annotated miRNAs (from 1,042 distinct loci) are never expressed at greater than 100 cpm in any cell or tissue, calling into question the likelihood they ever play endogenous roles as miRNAs. 1777 miRNAs (from 1268 loci) are never expressed above 500 cpm.

**Figure 3.**
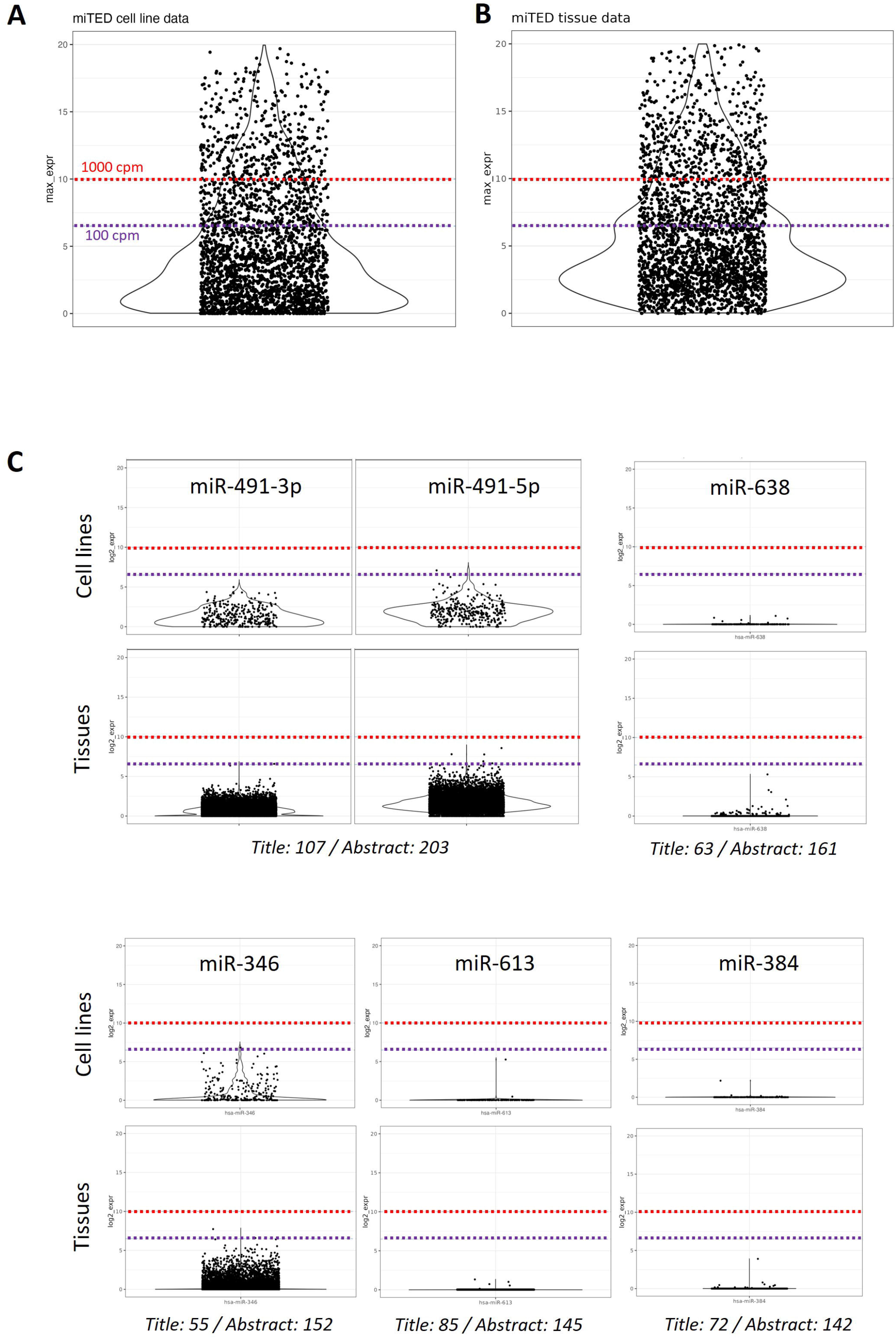
Assessment of maximal miRNA expression across cell lines and tissues. The maximal cpm expression for each of the 2656 miRBase annotated miRNAs is shown across all small RNA-seq data present in miTED (19): A) 333 cell lines and B) 10,740 tissues drawn from 176 different tissue sites. Log2 values are graphed with dotted lines indicating 100 and 1000 cpm. C) Expression for individual miRNAs across cell lines and tissues. Title / Abstract refers to the number of publications featuring that miRNA in the paper title or abstract by Pubmed search.

MicroRNAs expressed below the 100 cpm threshold are unlikely to be biologically important, however there are >4,500 publications where the name of a miRNA never expressed above 100 cpm features in the title, and >10,000 publications where it features in the abstract. Expression profiles are shown for the 5 most highly published, lowly expressed miRNAs (Figure 3C). miR-491-5p is also included to accompany miR-491-3p which itself is barely expressed above 100 cpm and only in a couple of samples.

Canonical miRNA biogenesis is a two step process, commencing in the nucleus when Drosha cleaves the base of an internally paired hairpin structure, then completed in the cytoplasm as Dicer cleaves both strands of the loop to liberate a small RNA duplex of which one strand is loaded into AGO. This strand is called the “guide” and serves as the functional mature miRNA, whilst the opposing strand (often called the “passenger”) remains un-incorporated and is degraded. For our analysis we chose to keep all 2,656 annotated miRNAs, some of which are established “guides”, others “passengers” (that might not be expected to play miRNA roles) and others where 5p and 3p arms are not formally annotated, in which case reads mapping across such loci are included in a single additive count. We chose to include all annotated miRNAs because some “passenger” arms may still be incorporated into RISC and function as miRNAs, whilst “arm-switching” may occur where the identity of the guide and passenger alters depending upon context. We note several extreme examples of this (Supplementary Figure 2) where extensively studied miRNAs vary dramatically in the proportions of 5p and 3p arms detected depending on the tissue (20). By keeping all miRNA annotations, regardless of whether or not they are assumed to be “passengers” or “guides”, and by drawing on data that is extensive in both cell/tissue types and numbers of samples, it is unlikely we would simply be missing cell contexts in which apparently lowly expressed miRNAs do in fact meet minimal expression thresholds where measureable activity could be expected.

### Some miRNAs show highly tissue-specific expression

In Figure 2 we observe that over half of annotated miRNAs are never expressed at a level whereby biologically detectable functions should be expected. In other instances, miRNAs are expressed at seemingly functional levels, however expression may be highly tissue/cell specific. We highlight three extreme examples; miR-567, miR-630 and miR-802, which are never detected at high levels in any tissue and which are only expressed highly in a single cell line (Figure 4A). However, even this appears to be a false positive as this is the same dataset for all three miRNAs, derived from HeLa cells, where independent HeLa cell data fails to detect their expression. Nonetheless, these same miRNAs have featured in 29, 125 and 111 publications respectively. Wider examination of >10k tissues, whose 176 different tissue annotations were curated into 54 basic tissue types, highlights striking tissue-specificity of dozens of well-studied miRNAs (Figure 4B), meaning that in most of these cases, “standard” cell lines (293, HeLa, A549 etc) will not represent a biologically appropriate context in which to examine function.

**Figure 4.**
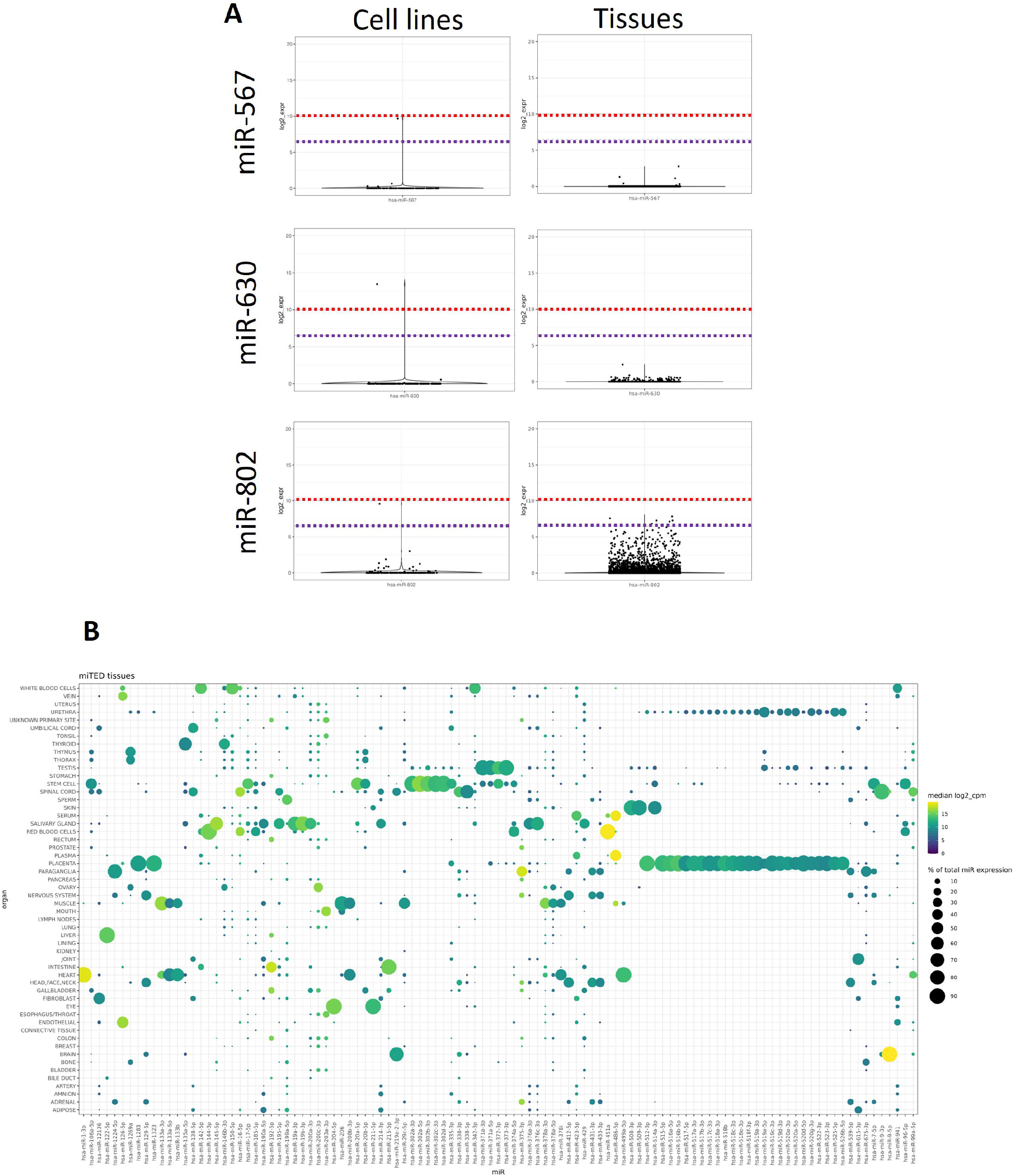
Tissue-restricted miRNA expression. A) miTED-derived maximal miRNA expression across cell lines and tissues for the miRNAs indicated. B) The 176 different annotated tissues in miTED were manually curated into 54 major tissue types (keeping all data), and the expression of the most tissue-specific miRNAs shown. Log2 cpm expression is indicated by colour and proportion of total miRNA expression for that miRNA is indicated by circle size. Only miRNAs expressed at greater than 500 cpm in at least one tissue are included in the plot.

## Discussion

In exactly the same way one can introduce synthetic siRNAs that don’t exist in nature to silence genes, one can introduce sequences representing miRNAs that will repress targets, even if the miRNA itself is barely detected in the endogenous context. Therefore, publications that are reliant upon miRNA over-expression can describe apparent phenotypic effects and demonstrate gene targeting, without ever establishing an endogenous role.

Based on our findings in the highly sensitive reporter system we have created, miRNAs that are present at less than 200 cpm (0.02% of total miRNA expression) do not show any evidence of detectable function. It is not that such miRNAs are inherently incapable of binding AGO, but rather the proportion to which they represent the gene targeting pool is so small their silencing, even of reporters that are far more sensitive than endogenous transcripts are simply undetectable. The surprise however is that for over half of all annotated miRNAs, their endogenous expression never reaches this level, which in turn calls into question the significance of multiple thousands of publications of which they are the collective subject.

Whilst it is impossible to state that for any given miRNA, there is no cellular context where it is ever highly expressed, the fact that our bioinformatic analysis involves >300 cell lines and >10,000 tissues from 176 different tissues and pathologies strongly supports our claim that the significance of most, if not all of these small RNAs, can be dismissed simply on the basis of expression alone, let alone ongoing debates about which small RNAs represent genuine miRNAs and which are misannotated products of the RNA degradome (6-8, 28-31).

Collectively, it is true that miRNAs en masse may exert “crowd control” where the summation of multiple small effects could add up to significant regulation (32), and it is established that the RISC component GW182 can organise multiple miRNA:AGO complexes to co-target a specific transcript which increases the strength of the targeting effect (33, 34). Even so, it is unlikely such mechanisms would make biologically significant, those miRNAs that are so lowly expressed that their repressive effects are undetectable in our highly sensitive reporters.

## Acknowledgements

This work was supported by funding from the Australian Research Council (FT190100544, DP190103333), the Cancer Council Beat Cancer Project (PRF2518) and the Worldwide Cancer Research Foundation (WCR-19-0300). K.A.P. was supported by the Royal Adelaide Hospital Research Committee Florey Fellowship. The authors would like to thank Prof Gregory Goodall for his critical reading of the manuscript.

## Author contributions

SAK performed the cloning and luciferase reporter assays. JMB led the bioinformatic work supported by KAP. PAG provided oversight, intellectual contribution and critical editing. CPB directed the project, led data analysis and wrote the manuscript.

