## Supplementary figures and images for "More than half of annotated human miRNAs are never expressed at levels sufficient for biological function"

### Supplemental Figure 1

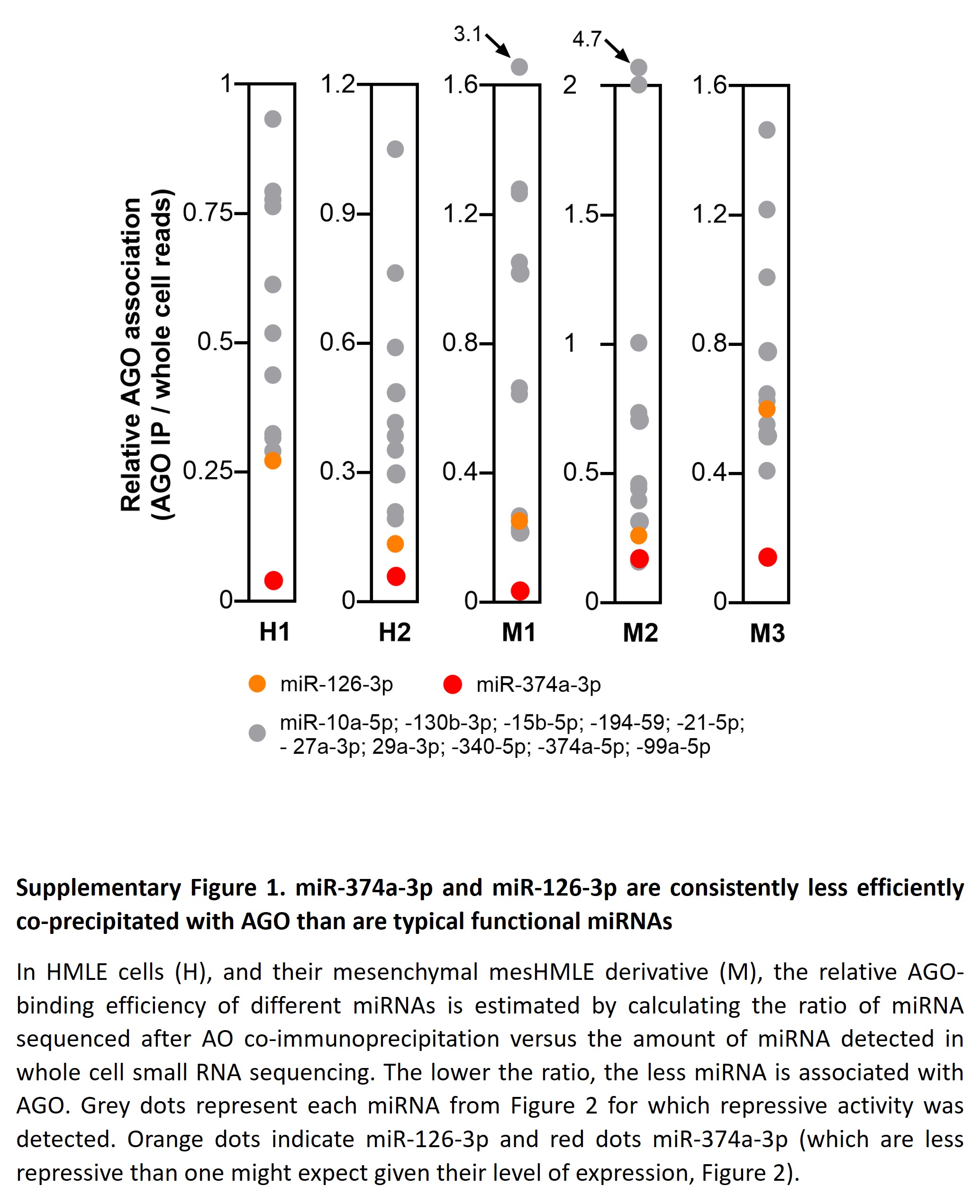

### Supplemental Figure 2

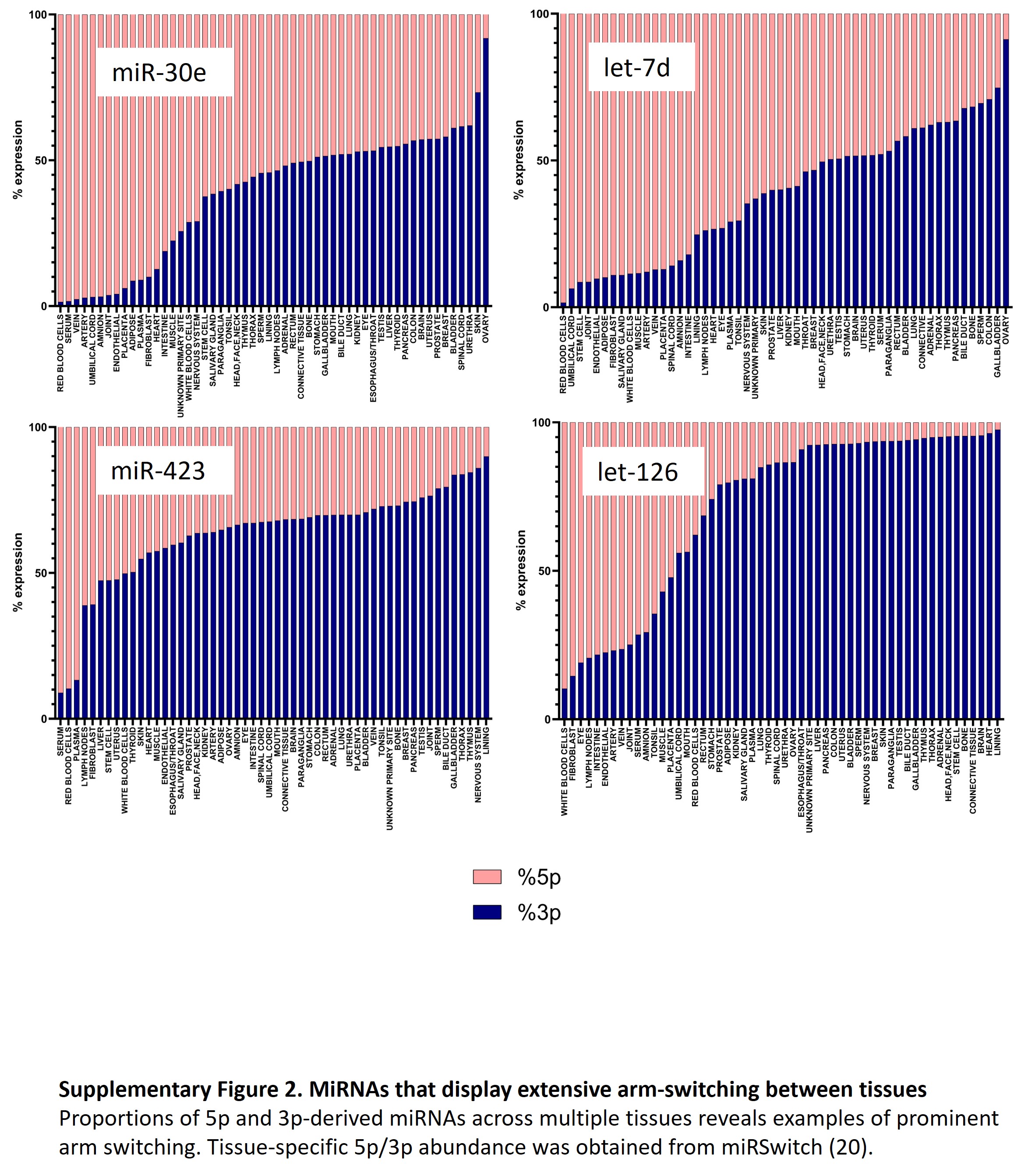
